# MEC-2/Stomatin is required for aversive behaviour but dispensable for prey detection in the predatory nematode *Pristionchus pacificus*

**DOI:** 10.64898/2026.03.09.710502

**Authors:** Marianne Roca, James W. Lightfoot

## Abstract

Sensory systems provide animals with essential information about their environment and are critical for generating appropriate behaviours. Mechanosensation is a fundamental component of this sensory repertoire, and disruption of mechanosensory pathways can have severe functional consequences. In the nematode *Caenorhabditis elegans*, mechanosensory circuits have been extensively characterized and mediate touch-driven navigation and avoidance. These circuits rely on conserved molecular components including the stomatin-like protein MEC-2 along with MEC-6, which function together in the mechanotransduction complex. In contrast, the predatory nematode *Pristionchus pacificus* has repurposed mechanosensory pathways to also enable prey detection, a derived ecological behaviour. As we previously demonstrated that *Ppa-mec-6* is required for efficient predation, here we assessed if *Ppa-mec-2* plays a similar role in *P. pacificus* prey detection. We find that while *Ppa-mec-2* is required for the aversive touch response, it is dispensable for prey detection. This functional divergence reflects differential neuronal expression as *Ppa-mec-2* is absent from the IL2 neurons that mediate prey detection and also robustly express *Ppa-mec-6*. These findings reveal that partitioning of mechanosensory components across neuronal types enables functional specialization, demonstrating how conserved sensory machinery can support distinct behavioural functions across evolution.

---

Sensory systems are fundamental to animal life, providing critical information about environmental conditions and physiological states. Mechanosensation enables organisms to detect touch, pressure, and noxious stimuli and these capabilities are essential for survival across diverse ecological contexts. The mechanosensory machinery is evolutionarily ancient and highly conserved, yet supports remarkably diverse functions (1). Examples of such diversity include its role mediating acoustic courtship through antennal mechanoreceptors in *Drosophila* (2), enabling flow-guided navigation and prey detection through the lateral line organs in fish (3), and facilitating hearing in vertebrates through hair-cell activation (4,5). Additionally, dysfunction in mammalian mechanosensory pathway’s may cause conditions ranging from chronic pain to complete sensory loss further underscoring its functional importance (6,7). Thus, understanding how conserved mechanosensory components are adapted to support different behaviours across species can illuminate both the evolutionary flexibility and functional constraints of sensory systems.

Fundamental discoveries in mechanotransduction have been made using the nematode *Caenorhabditis elegans*, a model organism renowned for its genetic tractability. In this nematode, mechanosensation influences egg laying, mating, feeding and locomotion (8). When exploring their environments, worms use mechanosensory information to navigate around obstacles (9) or escape danger like traps from predatory fungi (10). In the lab, *C. elegans* shows escape and avoidance behaviours upon exposure to vibrations or direct contact. These behaviours have been used to identify three main types of mechanosensation that differ by their sensitivity, and the neurons involved. These are harsh touch, related to nociception, as well as gentle touch and nose touch (11–13). Harsh touch is associated with the activity of more than 14 neurons including FLP, PVD, AQR, ADE, PDE, PHA/PHB among others. Gentle touch is characterised by the activity of 6 neurons such as ALMs, AVM, PLMs and PVM projecting along the body while nose touch triggers around 26 head neurons (including FLP, ASH, OLQ) (11–15). Similarly, the molecular mechanisms for each touch detection type only partially overlap (12–19) with the best characterised response associated with gentle touch. Genetic screens have led to the identification of numerous components associated with these processes including the *mec* (mechanosensory abnormal) genes (20). These include the Deg/ENaC components *Cel-mec-4* and *Cel-mec-10* that encode subunits of mechanosensitive ion channels in the touch receptor neurons of *C. elegans* (*16*), and form the basis of its response to gentle touch stimuli (17). In addition, several aspects of the *C. elegans* touch responses rely on the Transient Receptor Potential (TRP) family (8,13,21). Crucially, both the Deg/ENaC and TRP gene families are highly functionally conserved across diverse species (22,23).

Despite the functional conservation of mechanosensory pathways for detecting physical perturbations, different species utilise these sensory capabilities to support distinct behavioural outputs. Even among relatively simple organisms such as nematodes, mechanosensation serves diverse functions. While in *C. elegans*, touch primarily elicits a stereotyped aversive response, in the satellite model system *Pristionchus pacificus*, mechanosensory cues have recently been linked to not only an aversive response but also a broader repertoire of more complex behaviours (24). Specifically, *P. pacificus* utilises mechanosensation for predatory behaviour enabling it to detect prey and attack the larvae of other nematodes but not kin or close relatives (25–27). This predatory behaviour is dependent teeth-like denticles and a developmentally plastic mouth form with only one of two potential morphs capable of predation (28). In the predatory morph, aggressive behaviour is induced through the balance between the neuromodulators octopamine and tyramine which promote predatory or docile states respectively (29). In addition, serotonin coordinates the mobility and rhythmicity of one of the teeth to ensure successful predation (30). Importantly, prey detection depends on the integration of both chemosensory and mechanosensory inputs (31,32). While little is known of the importance of chemical cues and how they are recognised (33,34), several canonical mechanosensory components have been identified and have been co-opted for prey detection (12,31). Thus far, these consist of *Ppa-mec-3,* a transcription factor that promotes the expression of genes belonging to the mechanosensory pathway (35) and *Ppa-mec-6*, a paraoxonase-like protein, which enhances mechanosensitive channel function (36,37). However, as these represent regulatory and channel support components, they do not directly detect mechanical stimuli and transmit this information. Therefore, we further explored the molecular mechanisms allowing detection of mechanosensory stimuli in the context of predation and investigated other potential accessory mechanosensory channel proteins.

In *C. elegans*, *Cel-mec-2* is a stomatin-like protein that functions as an accessory protein in the mechanotransduction complex. Together with *Cel-mec-6,* they ensure the proper folding, trafficking and surface localisation of the channel subunits *Cel-mec-4* and *Cel-mec-10* (37,38). Furthermore, *mec-2* is the orthologue to the podocin encoding NPHS2 whose mutation leads to nephrotic syndrome in humans (39). Given the functional association in *C. elegans* between *Cel-mec-6* and *Cel-mec-2* and the additional predatory role of *Ppa-mec-6* in *P. pacificus*, we explored *Ppa-mec-2* for evidence of involvement in the evolution of prey detection in *P. pacificus*. Here, we show that mutations of *Ppa-mec-2* disrupt touch perception and reduce exploratory behaviour. However, we find that *Ppa-mec-2* is not required for prey detection and predation. Finally, we observed *Ppa-mec-2* expression in a different set of head cells than *Ppa-mec-6* suggesting that the spatial regulation of mechanosensory genes is associated with their involvement in predation.

## Results

Predation in *P. pacificus* requires mechanosensation but thus far *Ppa-mec-6* is the only component associated with channel function that also has a clear role in prey detection (31). Therefore, we explored another core mechanosensory gene canonically involved as an accessory protein in the mechanotransduction complex in *C. elegans* for any role in prey detection (37,38). This is the *stomatin* subunit *mec-2.* The *Cel-mec-2* gene is located on the X chromosome and is predicted to produce 17 isoforms (40,41). This extensive transcript diversity is conserved in *P. pacificus,* where the orthologous gene (*Ppa-mec-2*) is instead located on chromosome I and is predicted to generate 23 isoforms (Fig. S1), (42). The large number of isoforms together with the complex exon structure made it difficult to induce a mutation to fully disrupt the gene function. However, in *C. elegans* the mechanosensory function of *Cel-mec-*2 was shown to require the isoform E (40,41). Therefore, we aligned the *Ppa-mec-2* translated sequence to the *Cel-mec-2E* isoform. We identified that the translation of the *Ppa-mec-2* fourth exon shared 100% identity with a part of *Cel-mec-2E* isoform. Additionally, this fourth exon appears to be retained in all 23 predicted *P. pacificus* isoforms (Fig. 1a, b and S1). Consequently, we selected this exon as the target site for CRISPR/Cas9 to study the role of MEC-2 in predation. We successfully generated two alleles each with a distinct four base deletion with both mutations leading to an early codon stop and are predicted null mutants (Fig. 1b and table S1).

**Figure 1.**
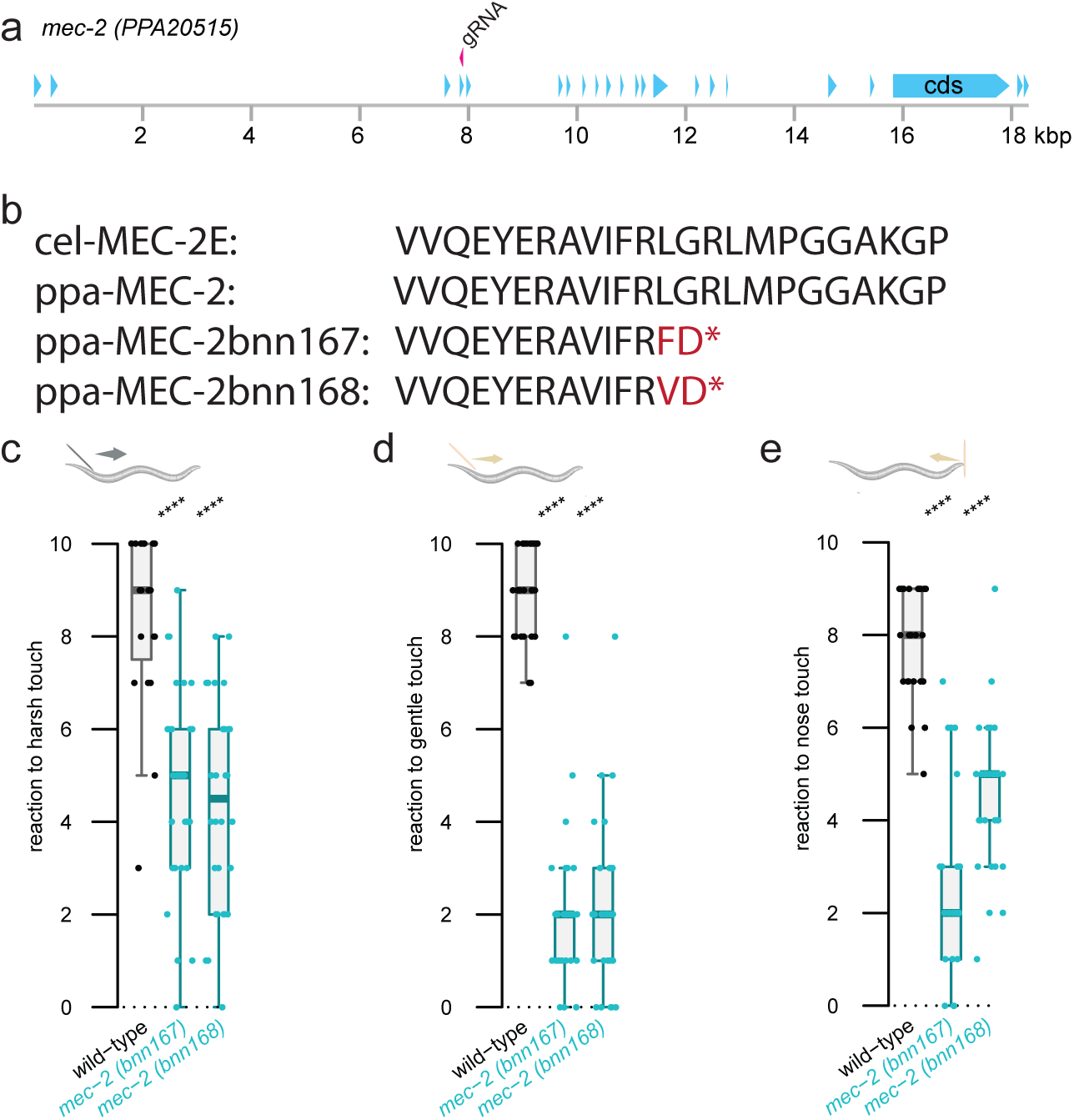
*Ppa-mec-2* is involved in touch detection. (A) Gene structure of *Ppa-mec-2*. The position of the gRNA use in this study is indicated in red. (B) Translated sequences of *Ppa-mec-2* fourth exon in wild-type and in the two mutant alleles generated in this study. The corresponding *Cel-mec-2E* sequence is also indicated. (C-E) Responses to the three main types of mechanosensation in touch assays: harsh touch (C), gentle touch (D) and nose touch (E). Each worm was assessed 10 times in a row and 30 worms were tested per strain. Statistical tests: two-tailed Wilcoxon Mann Whitney with Benjamin-Hochberg correction: non-significant (ns), p-value ≤ 0.05 (*), ≤ 0.01 (**), ≤ 0.001 (***), ≤ 0.0001 (****).

In *C. elegans,* mutations in *mec-2* disrupt the animals perception of gentle touch (37,41,43,44). Therefore, we first investigated if the mechanosensory role of *Ppa-mec-2* is conserved in *P. pacificus*. We assessed our *Ppa-mec-2* mutants using the three classical nematode touch assays. The worms’ sensitivity to harsh and gentle touch were determined through contact with the posterior body, and their response to nose touch by determining the number of responses to nose contact events (11,31). In both *Ppa-mec-2* alleles, we observed a strong reduction in their reaction to harsh body and gentle touch as well as nose touch (Fig. 1c-e). These results indicate that mutations in the fourth exon of *Ppa-mec-2* disrupt protein function and induce mechanosensory defects. Additionally, this confirms that the mechanotransduction role of this gene in contact-induced avoidance behaviour is conserved in *P. pacificus*.

Mechanosensation is not the only sensory system that may depend on *Ppa-mec-*2. Indeed, in *C. elegans* expression of *Cel-mec-2* in head neurons has also been linked to chemosensory functions (*40,41*). To explore this in *P. pacificus*, we performed chemotaxis assays to test avoidance toward the known *P. pacificus* repellent 1-octanol (31). We observed strong avoidance behaviour in both wild-type and *Ppa-mec-2* mutants indicating unchanged chemosensory function in *Ppa-mec-2* mutants (Fig. S2a). In *C. elegans*, chemosensory defects only occur from mutations in specific *Cel-mec-2* isoforms, but not for example in mutants in isoform MEC-2E. (*40,41*). Thus, while the mutated exon in our *Ppa-mec-2* mutants theoretically disrupts all isoforms, further analysis will be necessary to rule out any chemosensory function for *Ppa-mec-2*.

A striking characteristic of *P. pacificus* is its developmental plasticity which produces two alternative mouth morphs. The eu predatory morph is identified by its wide mouth with two teeth. In contrast the non-predatory st animals have a narrower mouth with only a single tooth (25). Several genetic and environmental factors influence the developmental choice, however, under our laboratory conditions the wild-type strain PS312 used throughout this study is highly eu (Fig. S2b) (28). Similarly, we noticed a high occurrence of the predatory mouth form eu in both *Ppa-mec-2* mutants (Fig. S2b). Therefore, similar to other touch mutants (31), *Ppa-mec-2* does not influence the mouth form developmental decision.

As both *Ppa-mec-2* mutants acquired a predatory eu morph, we next tested the role of *Ppa-mec-2* in prey detection. We assessed if the efficiency of predation was disrupted in our mutants by performing corpse assays (25). As previously, we used *C. elegans* larvae as potential prey and introduced five *P. pacificus* adults as potential predators. After 2 hours, the number of corpses left on the plate allowed us to evaluate killing efficiency (Fig. 2a) (25,31). When performing this assay with the mechanosensory deficient *Ppa-mec-6* mutants, the corpse number was reduced to approximately half of wild-type numbers (31). However, we did not observe any significant reduction in the number of corpses in either *Ppa-mec-2* mutant suggesting that the mutants were still efficiently finding prey (Fig. 2b). Instead, we observed that corpses generated by *Ppa-mec-2* mutants were more spatially clustered than those produced by wild-type animals, indicating that predatory events occur more locally in the mutants. To investigate this further, we quantified locomotion and exploration using an automated behavioural tracking system (29). Compared with wild-type animals, *Ppa-mec-2* mutants exhibited markedly reduced exploration (Fig. 2c), suggesting that *Ppa-mec-2* dependent mechanosensation promotes exploratory behaviour but is not required for predation itself.

**Figure 2.**
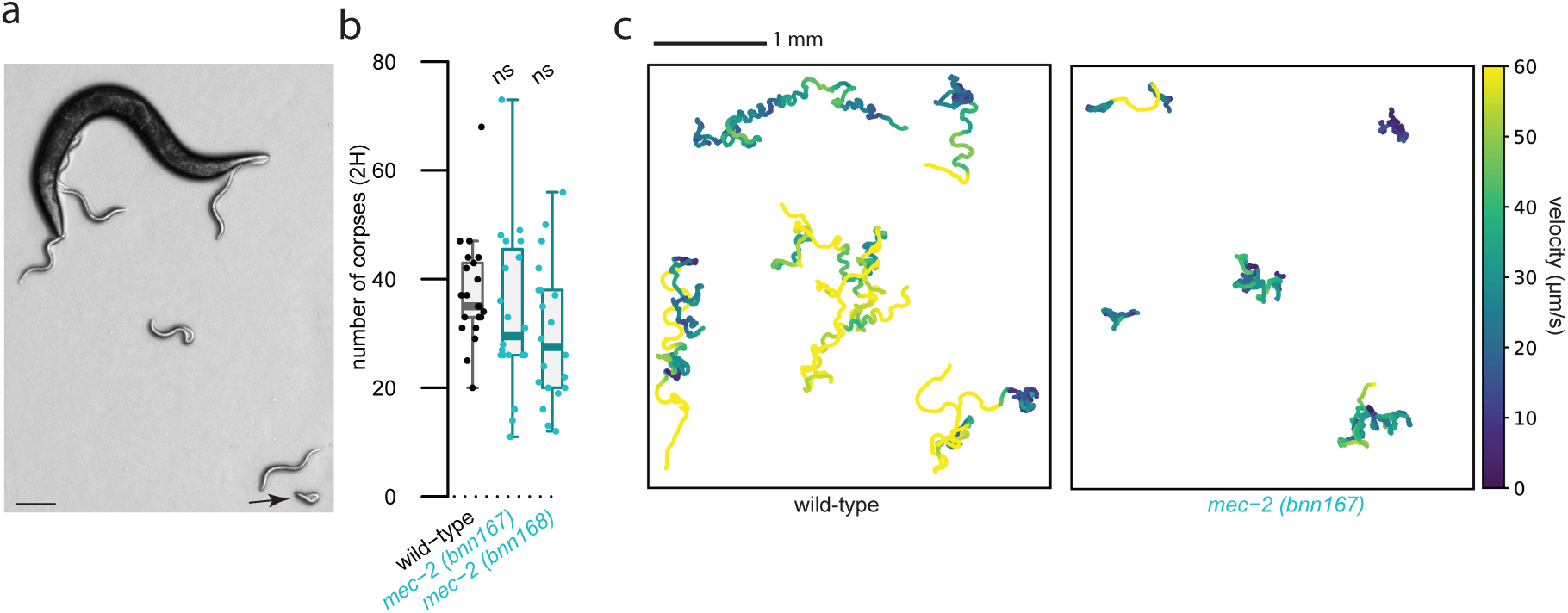
*Ppa-mec-2* mutants are predatory but have exploratory defects. (A) *P. pacificus* adult predation on *C. elegans* larvae. The arrow indicates a corpse. Scale bar is 100 μm. (B) Results of corpses assay with wild-type or two different *Ppa-mec-2* mutants as predators. The assay was repeated 20 times per strain. Statistical tests: two-tailed Wilcoxon Mann Whitney with Benjamin-Hochberg correction: non-significant (ns), p-value ≤ 0.05 (*), ≤ 0.01 (**), ≤ 0.001 (***), ≤ 0.0001 (****). (C) Locomotion of wildtype compared to *Ppa-mec-*2 mutants. Five representative trajectories of control and *Ppa-mec-2* mutant worms recorded in the presence of *C. elegans* prey are shown. The colour code gives the velocity (μm/s) at each position.

MEC-2 and MEC-6 are well established core components of the gentle touch mechanotransduction complex in touch receptor neurons. In this context both MEC-2 and MEC-6 are required for proper channel function and loss of either disrupts mechanosensation (37,38). However, as only mutations in *Ppa-mec-6* disrupts prey detection in *P. pacificus*, we hypothesised that this functional divergence might be linked to differences in their expression and subsequent role in the specific neurons underlying prey sensing and touch avoidance. In *P. pacificus, Ppa-mec-6* is expressed in several sensory neurons including robust expression in the six IL2 head sensory cells which function as part of the prey detection circuit (31). In addition, the IL2 neurons also express octopamine receptors that regulate predatory aggression in *P. pacificus* indicating both prey detection and aggression combine in the same sensory circuitry (29). In *C. elegans* the six environmentally exposed polymodal sensory IL2 neurons can stimulate locomotion under specific conditions (45) and are canonically involved in nictation behaviour. This is a behaviour frequently observed in dauer allowing dispersal of this environmentally resistant larval stage. The role of IL2 neurons in nictation was postulated to rely on stage specific dendritic arborizations in the dauer as well as in their perception of mechanosensory signals (46–48). Importantly, these neurons display species-specific morphologies. While both species extend anteriorly directed neurites with exposed termini, the IL2D and IL2V soma are located substantially more anteriorly in *P. pacificus* than in *C. elegans* (49). Furthermore, robust expression of *Ppa-mec-6* in these neurons is thought to be essential for their role in prey detection and, silencing these neurons also induces prey detection defects (29,31). Therefore, we decided to explore the expression patterns of *Ppa-mec-2* to investigate its potential presence in the IL2 neurons. As *Ppa-mec-2* has multiple isoforms, we could not rely on transcriptional reporters that are frequently used in *P. pacificus* studies (50). As such we instead used Hybridization Chain Reaction (HCR). This is a variant of smFISH that uses enzyme-free signal amplification, whereby initiator probes bind to target transcripts triggering the polymerization of fluorescent DNA hairpins to enhance detection sensitivity in situ. This was recently successfully established in *P. pacificus* (51,52). Using this method allowed us to cover all potential exon sequences with probes without requiring prior knowledge of cis-regulatory element localisation. Consistent with previous observations using a reporter line (31), *Ppa-mec-6* showed the strongest signals in head cells whose positions correspond to the IL2 neurons (Fig. 3a). In contrast, *Ppa-mec-2* expression was mainly detected in a few cells along the body (Fig. 3b), most likely corresponding to the 6 touch receptor neurons AVM, ALMs, PVM, PLMs known to express *Cel-mec-2* in *C. elegans* (*40,41*). Additionally, weak *Ppa-mec-2* signal was observed in the head as previously described in *C. elegans* (*43*). However, head cells expressing *Ppa-mec-2* were distinct from those expressing *Ppa-mec-6* (Fig. 3c). This discrepancy in expression patterns likely underlies the functional divergence between the two genes.

**Figure 3.**
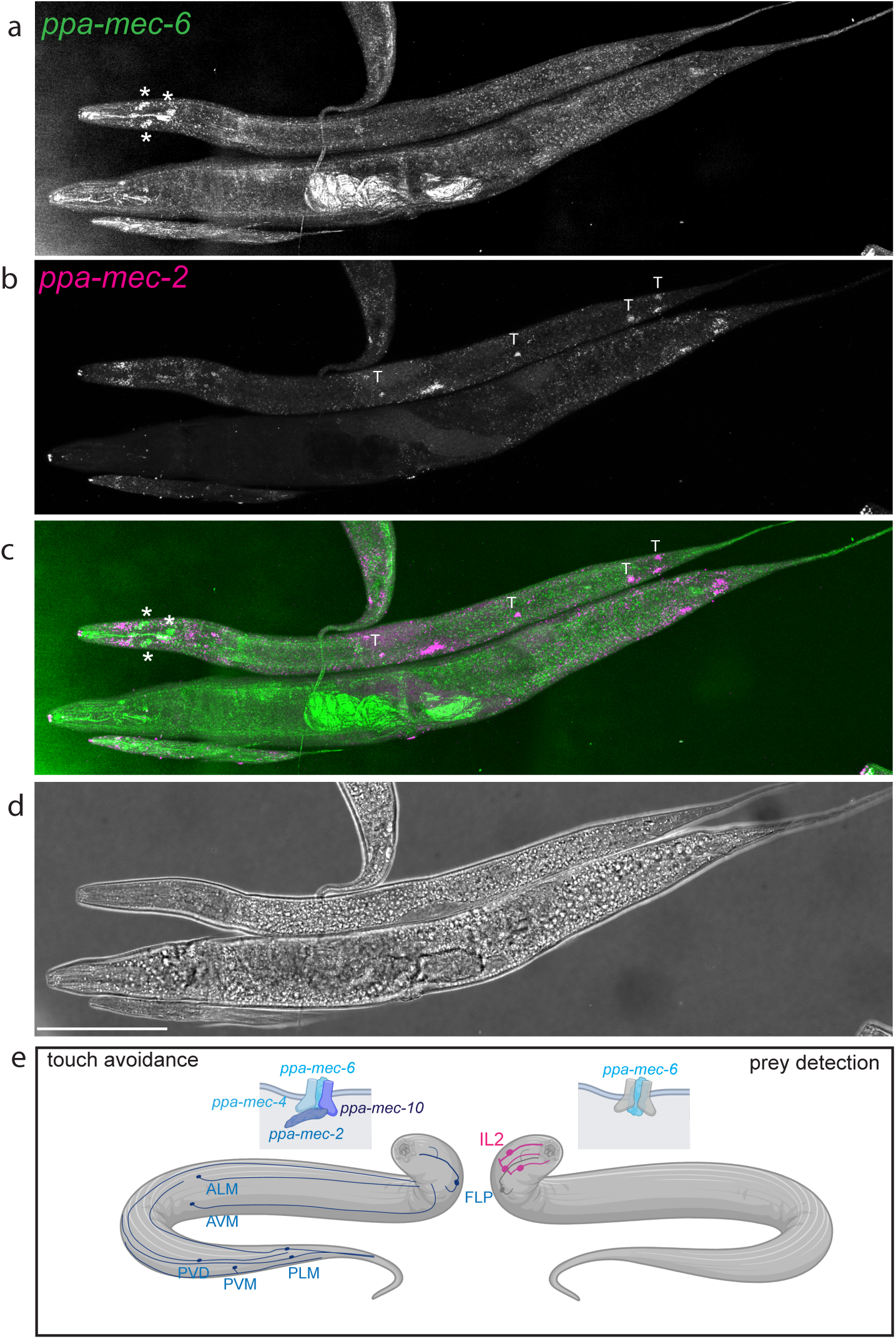
*Ppa-mec-2* is not expressed in neurons known to mediate prey detection. Representative images of worms stained with HCR for (A) *Ppa-mec-6* and (B) *Ppa-mec-2*. (C) Merged image of *Ppa-mec-6* (green) and *Ppa-mec-2* (magenta). IL2 (*) and putative touch neurons (T) are indicated. Note that in the merge signals for both transcripts do not co-localize. (D) Brightfield of stained worms. Scale bar is 100 μm. (e) In *P. pacificus*, two mechanosensory pathways exist: one facilitates touch avoidance (left), while the other mediates prey detection (right).

## Discussion

In this study, we took advantage of behavioural variation between the nematodes *P. pacificus* and *C. elegans* to explore mechanistic changes during the co-evolution of mechanosensation and predation. Our results hint at a modular organisation of sensory pathways, in which individual components can be assembled in distinct combinations depending on cellular context and behavioural requirements.

Combining results from this study alongside previous data (7), we demonstrated that a mechanosensory pathway relying on both *Ppa-mec-6* and *Ppa-mec-2* as well as the channels *Ppa-mec-4* and *Ppa-mec-10* enables worms to detect and avoid potentially harmful contact (Fig. 3e). Our detailed behavioural analyses further revealed a marked reduction in exploratory activity in *Ppa-mec-2* mutants, particularly in high-velocity movements. This reduction resembles, but is more pronounced than the phenotype previously observed in *Ppa-mec-6* mutants (7). Moreover, similar lethargic behaviour has been repeatedly reported in *mec* mutants of *C. elegans* and is thought to reflect a cautious behavioural state in animals with impaired sensory input (53). Importantly, both *Ppa-mec-6* and *Ppa-mec-2* are expressed in several putative touch receptor neurons where they are also observed in *C. elegans*. Therefore, this highlights that key elements of the canonical mechanosensory pathway are conserved at both the molecular and neuronal level (Fig. 3e) contributing to touch avoidance and exploratory behaviour.

Crucially, in *P. pacificus* specific mechanosensory components underlie the emergence of its predatory behaviour (Fig. 3e). Previously, we found that the co-option of *Ppa-mec-6* is required for efficient prey detection but this is not accompanied by the recruitment of *Ppa-mec-4* and *Ppa-mec-10* (*31*) nor as we demonstrated here, *Ppa-mec-2.* The functional divergence between *Ppa-mec-2* and *Ppa-mec-6* is likely attributed to differences in their spatial expression. In particular, *Ppa-mec-2* transcripts were absent from the IL2 head neurons that express *Ppa-mec-6* and mediate prey detection. Our findings underscore the importance of gene regulation in shaping sensory system function and reveal that *Ppa-mec-6* can operate independently of *Ppa-mec-2*. However, we suspect that not only the transcriptional control of *Ppa-mec-2* but additional regulatory mechanisms may enable the dual functionality of the mechanosensory pathway in *P. pacificus*. A similar strategy operates in mammals where the *mec-2 stomatin* orthologue has different regulatory effects on the activity of channels depending on the ASIC subunit with which it interacts (54). Consistent with this, regulatory changes associated with the co-option of core mechanosensory components into distinct neuronal circuits likely facilitated prey detection and the acquisition of predatory behaviours in *P. pacificus* (Fig. 3e).

The functional divergence between *Ppa-mec-2* and *Ppa-mec-6* in predation raises pertinent questions regarding the extent to which these genes roles differ and how these roles were acquired. In *C. elegans, Cel-mec-6* is primarily studied in the context of gentle touch where together with *Cel-mec-2* it enables the functionality of the *Cel-mec-4* and *Cel-mec-10* channels (12,36,37). In contrast, *Cel-mec-2* also functions in other neurons and sensory modalities. Notably, specific *Cel-mec-2* isoforms are implicated in chemosensation without apparent *Cel-mec-6* involvement. This role would be consistent with the broader sensory function reported in mammalian stomatin domain genes (40,41,55). While our data suggest that *Ppa-mec-2* is not required for chemosensory processes, further analysis will be necessary to determine any specific roles associated with distinct isoforms in *P. pacificus*. Similarly, as *Ppa-mec-2* is not detected in the IL2 neurons where *Ppa-mec-6* is robustly expressed and involved in prey detection, another stomatin domain gene may have assumed this mechanosensory role typically associated with *mec-2*. Dissecting these alternative configurations will be essential to understand how conserved mechanosensory components are differentially deployed across species to generate behavioural diversity.

Together, these findings support a view that mechanosensory pathways are evolutionarily modular, allowing individual components to be reassigned or replaced to support novel behavioural functions. The characterisation of this non-canonical mechanosensory pathway in *P. pacificus* underscores the value of studying sensory systems across diverse species and ecological contexts. By revealing how conserved molecular components and neuronal circuits can be differentially utilised, our work illustrates how variation in genetic pathway organisation can have profound physiological and behavioural consequences. More broadly, these insights highlight how evolutionary modifications to sensory systems contribute to behavioural diversification.

## Material and method

### Worm maintenance

The *P. pacificus* strains used in this study are derived from the PS312 background and are listed in supplementary table 1. Additionally, we used the *C. elegans* N2 strain provided by the CGC funded by NIH Office of Research Infrastructure Programs (P40 OD010440). All worms were grown at 20°C on 6 cm plates with nematode growth media (NGM) seeded with 300 μl of *Escherichia coli* (OP50).

### Mutagenesis

The *Ppa-mec-*2 gene was identified from the *Pristionchus pacificus* genome as PPA20515 and isoform sequences as Iso_D.3007.1:23 (56). Gene structures were drawn using both ‘benchling’ and ‘chatGPT-5’ using the information retrieved from Pristionchus.org. More specifically, predicted isoform sequences were aligned to the genomic sequence using a k-mer–based seed-and-extend strategy (minimum seed length 15 bp). Exon–intron structures were inferred from contiguous exact matches between transcripts and the genome, allowing for multiple non-contiguous segments per isoform (representing exons). Exon start and end coordinates (1-based, inclusive) were extracted for each isoform and tabulated. Gene models were visualised as exon–intron schematics, drawn to scale with exons represented as filled boxes and introns as connecting lines/spaces. The *Cel-mec-2E* sequence was retrieve from wormbase (57). Alignment of predicted translation was made using clustal omega (58). gRNA was manually designed and the sequence is available in supplementary table1 along with primer sequences. CRISPR/cas9 directed mutagenesis was conducted as previously described (59) using *Ppa-prl-1* as a marker. Co-induce *Ppa-prl-1* mutations were subsequently outcrossed. Sequences of the mutation is displayed in supplementary table 1.

### Behavioural assays

#### Touch assays

Touch assays were conducted as previously described (31). In brief, worms were touched on their posterior body with a metallic worm pick (harsh) or eyelash (gentle) or on the nose with an eyelash (nose) and any response assessed. Each worm was assessed 10 times in a row and 30 worms were tested per strain.

#### Chemosensory assays

Chemosensory assays tested avoidance of 1-octanol (31). To do so, the midline of a 6 cm assay plate was first marked. Subsequently, at 1.5 cm either side of the midline, a drop of sodium Azide (1.2 μl, 1M) was added. This was followed by a drop of 100 % ethanol on one side and 100 % 1-octanol (sigma-aldrich) on the other. Next, 3 h starved worms were then placed on the midline of a plate and after 16 hours at 20°C, the number of worms in a 2 cm radius of each drop was assessed and the ratio calculated to give the chemosensory index. Only plates with a minimum of 50 worms in the vicinity of either drop were assessed. Strains were tested on at least two independent days.

#### Corpse assays

Corpse assays were performed as in our previous study (31). Briefly, 5 *P. pacificus* adults previously starved for 2 hours were added to 1.5 μl of filtered (20 μm, Millipore) *C. elegans* larvae spread on an unseeded NGM plate in a 2.25 cm^2^ copper arena. After 2 additional hours, the arena was imaged twice a few seconds apart at 20x (Axio Zoom V16; Zeiss). Using Fiji/imageJ, the observable differences between the two images allowed larva carcasses to be distinguished from moving larvae.

#### Locomotion tracking

The tracking of worms to allow behavioural analysis was performed using our previously developed workflow (29). The strains used all carryied the *myo-2p*::RFP reporter (31) and velocity was calculated by tracking the movement of the pharyngeal fluorescence. We recorded at 30 fps for 10 minutes at 1x effective magnification (60 N-C ⅔’’ 0.5 x, Zeiss, acA3088-57um, BASLER, Axio Zoom V16, Zeiss). We adjusted the exposure time to prevent saturating the signal. Recordings were analysed using PharagGow (60) adapted for *P. pacificus* (29). Recordings were made on at least 3 different days for a total of at least 130 worms.

### Determination of mouth form

Worms were grown at 20°C on NGM seeded plates for multiple generations and until a high number of adults worm were present on the plate. Worms were then wash out in M9 and a transferred in a drop to a slide with 0.3M of sodium azide to aid observation. A minimum of 30 animals were scored per assay under an Axiovert 200 (zeiss) at 10x or 20x to ensure visual identification of eu or st morphological features. The assessment was performed five time for each strains using plates from at least three different days.

### Data representation and statistical analysis

Results are displayed as boxplots showing the first and third quartiles as the box, with a line marking the median. Whiskers delimit the spread of the data excluding outliers. Consistent with previous studies (31), two-direction Wilcoxon Mann Whitney with Benjamini-Hochberg correction was used and results are indicated as follows: non-significant (ns), p-value ≤ 0.05 (*), ≤ 0.01 (**), ≤ 0.001 (***), ≤ 0.0001 (****). Graphics were generated and statistics performed with R (61).

### Expression pattern with in situ hybridization chain reaction (HCR)

HCR was performed as described previously with probes designed using the provided script and available in supplementary table 2 (52). Worms of various stages from nearly starved plates were used. To successfully digest the cuticle, worms were soaked in proteinase K (>600U/ml, thermos scientist) for 1 h at 37°C. For the hybridization we used 5 μl of B Probe (IDT) at 50 pmol with *Ppa-mec-6* targeted by B3 (AF 488) and *Ppa-mec-2* by B3 (AF 488). Corresponding hairpin and buffer were purchased from Molecular Instrument. 1 μl of DAPI at 1 g/l was added during the penultimate wash. Worms were mounted in Prolong gold antifade reagent (Cell Signaling Technology) and imaged using a confocal stellaris microscope (Leica) with a 40x water objective. The experiment was performed two times with a large number of worms to ensure consistency.

## Figure legends

**Supplementary Figure 1: *Ppa-mec-2* isoforms**

*Ppa-mec-2* gene structure is displayed with coding sequences (CDS) in blue and the position of the gRNA indicated in red. Isoforms are shown below with exons structure in orange.

**Supplementary Figure 2: Chemosensation and mouth form ratio are not affected in *Ppa-mec-2***

(A) Chemosensory index toward 1-octanol is plotted for wild-type and a *Ppa-mec-2* mutant. The assay was performed five time per strain. (B) Percentage of eurystomatous (eu) worms was assessed five time per strain. Statistical tests: two-tailed Wilcoxon Mann Whitney with Benjamin-Hochberg correction: non-significant (ns), p-value ≤ 0.05 (*), ≤ 0.01 (**), ≤ 0.001 (***), ≤ 0.0001 (****).

**Supplementary Table 1: Strain information**

**Supplementary Table 2: HCR probe information**

## Acknowledgments

We would like to thank Dr Lewis A. Cockram for useful discussions and the Max-Planck-Institute for Neurobiology of Behavior – caesar microscope facility for imaging assistance. Schematic were made with Biorender.

## Data availability

All data are available in the main text or the supplementary. Raw data from behavioral tracking will be made available via file transfer upon request.

## Author Contributions

MR, JWL Conceptualization, MR methodology, MR investigation MR, JWL original draft preparation, MR, JWL writing, review and editing.

## Reference

1. Schüler A, Schmitz G, Reft A, Özbek S, Thurm U, Bornberg-Bauer E. The Rise and Fall of TRP-N, an Ancient Family of Mechanogated Ion Channels, in Metazoa. Genome Biol Evol. 22 juin 2015;7(6):1713-27. doi:10.1093/gbe/evv091 PubMed PMID: 26100409; PubMed Central PMCID: PMC4494053.

2. Effertz T, Wiek R, Göpfert MC. NompC TRP Channel Is Essential for *Drosophila* Sound Receptor Function. Current Biology. 12 avr 2011;21(7):592-7. doi:10.1016/j.cub.2011.02.048

3. Montgomery JC, Baker CF, Carton AG. The lateral line can mediate rheotaxis in fish. Nature. oct 1997;389(6654):960-3. doi:10.1038/40135

4. Corey DP, Hudspeth AJ. Ionic basis of the receptor potential in a vertebrate hair cell. Nature. oct 1979;281(5733):675-7. doi:10.1038/281675a0

5. Pan B, Géléoc GS, Asai Y, Horwitz GC, Kurima K, Ishikawa K, et al. TMC1 and TMC2 are components of the mechanotransduction channel in hair cells of the mammalian inner ear. Neuron. 7 août 2013;79(3):504-15. doi:10.1016/j.neuron.2013.06.019 PubMed PMID: 23871232; PubMed Central PMCID: PMC3827726.

6. Baron R, Binder A, Wasner G. Neuropathic pain: diagnosis, pathophysiological mechanisms, and treatment. The Lancet Neurology. 1 août 2010;9(8):807-19. doi:10.1016/S1474-4422(10)70143-5

7. Rotthier A, Baets J, Vriendt ED, Jacobs A, Auer-Grumbach M, Lévy N, et al. Genes for hereditary sensory and autonomic neuropathies: a genotype–phenotype correlation. Brain. oct 2009;132(10):2699-711. doi:10.1093/brain/awp198 PubMed PMID: 19651702; PubMed Central PMCID: PMC2759337.

8. Bounoutas A, Chalfie M. Touch sensitivity in Caenorhabditis elegans. Pflugers Arch - Eur J Physiol. 1 août 2007;454(5):691-702. doi:10.1007/s00424-006-0187-x

9. Nam SW, Qian C, Kim SH, van Noort D, Chiam KH, Park S. C. elegans sensing of and entrainment along obstacles require different neurons at different body locations. Sci Rep. 27 nov 2013;3:3247. doi:10.1038/srep03247 PubMed PMID: 24284409; PubMed Central PMCID: PMC3842086.

10. Maguire SM, Clark CM, Nunnari J, Pirri JK, Alkema MJ. The C. elegans touch response facilitates escape from predacious fungi. Curr Biol. 9 août 2011;21(15):1326-30. doi:10.1016/j.cub.2011.06.063 PubMed PMID: 21802299; PubMed Central PMCID: PMC3266163.

11. Chalfie M, Hart AC, Rankin CH, Goodman MB. Assaying mechanosensation. WormBook. 31 juill 2014. doi:10.1895/wormbook.1.172.1 PubMed PMID: 25093996; PubMed Central PMCID: PMC4448936.

12. Goodman M. Mechanosensation. WormBook. 2006. doi:10.1895/wormbook.1.62.1

13. Li W, Kang L, Piggott BJ, Feng Z, Shawn Xu XZ. The neural circuits and sensory channels mediating harsh touch sensation in C. elegans. Nat Commun. mai 2011;2:315. doi:10.1038/ncomms1308 PubMed PMID: 21587232; PubMed Central PMCID: PMC3098610.

14. Goodman MB, Sengupta P. How Caenorhabditis elegans Senses Mechanical Stress, Temperature, and Other Physical Stimuli. Genetics. mai 2019;212(1):25-51. doi:10.1534/genetics.118.300241 PubMed PMID: 31053616; PubMed Central PMCID: PMC6499529.

15. Schafer WR. Mechanosensory molecules and circuits in C. elegans. Pflugers Arch. 2015;467(1):39-48. doi:10.1007/s00424-014-1574-3 PubMed PMID: 25053538; PubMed Central PMCID: PMC4281349.

16. O’Hagan R, Chalfie M, Goodman MB. The MEC-4 DEG/ENaC channel of Caenorhabditis elegans touch receptor neurons transduces mechanical signals. Nat Neurosci. Janv 2005;8(1):43-50. doi:10.1038/nn1362

17. Árnadóttir J, O’Hagan R, Chen Y, Goodman MB, Chalfie M. The DEG/ENaC Protein MEC-10 Regulates the Transduction Channel Complex in Caenorhabditis elegans Touch Receptor Neurons. J Neurosci. 31 août 2011;31(35):12695-704. doi:10.1523/JNEUROSCI.4580-10.2011 PubMed PMID: 21880930; PubMed Central PMCID: PMC3172708.

18. Chatzigeorgiou M, Schafer WR. Lateral facilitation between primary mechanosensory neurons controls nose touch perception in C. elegans. Neuron. 28 avr 2011;70(2):299-309. doi:10.1016/j.neuron.2011.02.046 PubMed PMID: 21521615; PubMed Central PMCID: PMC3145979.

19. Han L, Wang Y, Sangaletti R, D’Urso G, Lu Y, Shaham S, et al. Two novel DEG/ENaC channel subunits expressed in glia are needed for nose-touch sensitivity in Caenorhabditis elegans. J Neurosci. 16 janv 2013;33(3):936-49. doi:10.1523/JNEUROSCI.2749-12.2013 PubMed PMID: 23325233; PubMed Central PMCID: PMC3711640.

20. Chalfie M, Sulston J. Developmental genetics of the mechanosensory neurons of *Caenorhabditis elegans*. Developmental Biology. 1 mars 1981;82(2):358-70. doi:10.1016/0012-1606(81)90459-0

21. Kindt KS, Viswanath V, Macpherson L, Quast K, Hu H, Patapoutian A, et al. Caenorhabditis elegans TRPA-1 functions in mechanosensation. Nat Neurosci. mai 2007;10(5):5. doi:10.1038/nn1886

22. Ben-Shahar Y. Sensory Functions for Degenerin/Epithelial Sodium Channels (DEG/ENaC). Adv Genet. 2011;76:1-26. doi:10.1016/B978-0-12-386481-9.00001-8 PubMed PMID: 22099690; PubMed Central PMCID: PMC3298668.

23. Árnadóttir J, Chalfie M. Eukaryotic Mechanosensitive Channels. Annual Review of Biophysics. 9 juin 2010;39(Volume 39, 2010):111-37. doi:10.1146/annurev.biophys.37.032807.125836

24. Sommer RJ, Lightfoot JW. The Genus Pristionchus: a Model for Phenotypic Plasticity, Predatory Behavior, Self-Recognition and Other Complex Traits. In: Nematodes as Model Organisms [Internet]. 2022 [cité 8 janv 2026]. p. 1-23. (CABI Books). Disponible sur: https://www.cabidigitallibrary.org/doi/10.1079/9781789248814.0001 doi:10.1079/9781789248814.0001

25. Wilecki M, Lightfoot JW, Susoy V, Sommer RJ. Predatory feeding behaviour in Pristionchus nematodes is dependent on phenotypic plasticity and induced by serotonin. J Exp Biol. Mai 2015;218(Pt 9):1306-13. doi:10.1242/jeb.118620 PubMed PMID: 25767144.

26. Lightfoot JW, Wilecki M, Rödelsperger C, Moreno E, Susoy V, Witte H, et al. Small peptide-mediated self-recognition prevents cannibalism in predatory nematodes. Science. 5 avr 2019;364(6435):86-9. doi:10.1126/science.aav9856 PubMed PMID: 30948551.

27. Lightfoot JW, Dardiry M, Kalirad A, Giaimo S, Eberhardt G, Witte H, et al. Sex or cannibalism: Polyphenism and kin recognition control social action strategies in nematodes. Sci Adv. août 2021;7(35):eabg8042. doi:10.1126/sciadv.abg8042 PubMed PMID: 34433565; PubMed Central PMCID: PMC8386922.

28. Ragsdale EJ, Müller MR, Rödelsperger C, Sommer RJ. A developmental switch coupled to the evolution of plasticity acts through a sulfatase. Cell. 7 nov 2013;155(4):922-33. doi:10.1016/j.cell.2013.09.054 PubMed PMID: 24209628.

29. Eren GG, Böger L, Roca M, Hiramatsu F, Liu J, Alvarez L, et al. Predatory aggression evolved through adaptations to noradrenergic circuits. Nature. 21 janv 2026;1-10. doi:10.1038/s41586-025-10009-x

30. Okumura M, Wilecki M, Sommer RJ. Serotonin Drives Predatory Feeding Behavior via Synchronous Feeding Rhythms in the Nematode Pristionchus pacificus. G3 (Bethesda). 6 nov 2017;7(11):3745-55. doi:10.1534/g3.117.300263 PubMed PMID: 28903981; PubMed Central PMCID: PMC5677172.

31. Roca M, Göze Eren G, Böger L, Didenko O, Lo WS, Scholz M, et al. Evolution of sensory systems underlies the emergence of predatory feeding behaviors in nematodes. Proceedings of the National Academy of Sciences. 3 févr 2026;123(5):e2514172123. doi:10.1073/pnas.2514172123

32. Moreno E, Lightfoot JW, Lenuzzi M, Sommer RJ. Cilia drive developmental plasticity and are essential for efficient prey detection in predatory nematodes. Proc Biol Sci. 9 oct 2019;286(1912):20191089. doi:10.1098/rspb.2019.1089 PubMed PMID: 31575374; PubMed Central PMCID: PMC6790756.

33. Hiramatsu F, Goetting DL, Kotowska AM, Zorn N, Chauhan VM, Lightfoot JW. Contact-based kin discrimination is associated with specific surface lipids in the cannibalistic nematode Pristionchus pacificus [Internet]. bioRxiv; 2026 [cité 10 févr 2026]. p. 2026.01.13.699210. Disponible sur: https://www.biorxiv.org/content/10.64898/2026.01.13.699210v1 doi:10.64898/2026.01.13.699210

34. Kotowska AM, Hiramatsu F, Alexander MR, Scurr DJ, Lightfoot JW, Chauhan VM. Surface Lipids in Nematodes are Influenced by Development and Species-specific Adaptations. J Am Chem Soc. 26 févr 2025;147(8):6439-49. doi:10.1021/jacs.4c12519

35. Way JC, Chalfie M. mec-3, a homeobox-containing gene that specifies differentiation of the touch receptor neurons in C. elegans. Cell. 1 juill 1988;54(1):5-16. doi:10.1016/0092-8674(88)90174-2 PubMed PMID: 2898300.

36. Chelur DS, Ernstrom GG, Goodman MB, Yao CA, Chen L, O’ Hagan R, et al. The mechanosensory protein MEC-6 is a subunit of the C. elegans touch-cell degenerin channel. Nature. déc 2002;420(6916):669-73. doi:10.1038/nature01205

37. Brown AL, Liao Z, Goodman MB. MEC-2 and MEC-6 in the Caenorhabditis elegans Sensory Mechanotransduction Complex: Auxiliary Subunits that Enable Channel Activity. J Gen Physiol. juin 2008;131(6):605-16. doi:10.1085/jgp.200709910 PubMed PMID: 18504316; PubMed Central PMCID: PMC2391253.

38. Zhang S, Arnadottir J, Keller C, Caldwell GA, Yao CA, Chalfie M. MEC-2 Is Recruited to the Putative Mechanosensory Complex in *C. elegans* Touch Receptor Neurons through Its Stomatin-like Domain. Current Biology. 9 nov 2004;14(21):1888-96. doi:10.1016/j.cub.2004.10.030

39. Boute N, Gribouval O, Roselli S, Benessy F, Lee H, Fuchshuber A, et al. NPHS2, encoding the glomerular protein podocin, is mutated in autosomal recessive steroid-resistant nephrotic syndrome. Nat Genet. avr 2000;24(4):349-54. doi:10.1038/74166

40. Keszthelyi TM, Légrádi R, Pálya D, Köles T, Regős Á, Karancsiné Menyhárd D, et al. The MEC-2E isoform with a large C-terminal completely rescues the touch sensation defect of C. elegans. Sci Rep. 22 juill 2025;15:26606. doi:10.1038/s41598-025-10711-w PubMed PMID: 40695872; PubMed Central PMCID: PMC12284166.

41. Liang X, Calovich-Benne C, Norris A. Sensory neuron transcriptomes reveal complex neuron-specific function and regulation of mec-2/Stomatin splicing. Nucleic Acids Res. 8 déc 2021;50(5):2401-16. doi:10.1093/nar/gkab1134 PubMed PMID: 34875684; PubMed Central PMCID: PMC8934639.

42. Werner MS, Sieriebriennikov B, Prabh N, Loschko T, Lanz C, Sommer RJ. Young genes have distinct gene structure, epigenetic profiles, and transcriptional regulation. Genome Res. nov 2018;28(11):1675-87. doi:10.1101/gr.234872.118 PubMed PMID: 30232198; PubMed Central PMCID: PMC6211652.

43. Huang M, Gu G, Ferguson EL, Chalfie M. A stomatin-like protein necessary for mechanosensation in C. elegans. Nature. nov 1995;378(6554):292-5. doi:10.1038/378292a0

44. Suzuki H, Kerr R, Bianchi L, Frøkjær-Jensen C, Slone D, Xue J, et al. In Vivo Imaging of *C. elegans* Mechanosensory Neurons Demonstrates a Specific Role for the MEC-4 Channel in the Process of Gentle Touch Sensation. Neuron. 11 sept 2003;39(6):1005-17. doi:10.1016/j.neuron.2003.08.015

45. Johnson JR, Edwards MR, Davies H, Newman D, Holden W, Jenkins RE, et al. Ethanol Stimulates Locomotion via a Gαs-Signaling Pathway in IL2 Neurons in Caenorhabditis elegans. Genetics. nov 2017;207(3):1023-39. doi:10.1534/genetics.117.300119 PubMed PMID: 28951527; PubMed Central PMCID: PMC5676223.

46. Schroeder NE, Androwski RJ, Rashid A, Lee H, Lee J, Barr MM. Dauer-specific dendrite arborization in C. elegans is regulated by KPC-1/Furin. Curr Biol. 19 août 2013;23(16):1527-35. doi:10.1016/j.cub.2013.06.058 PubMed PMID: 23932402; PubMed Central PMCID: PMC4671503.

47. Lee H, Choi M kyu, Lee D, Kim H sung, Hwang H, Kim H, et al. Nictation, a dispersal behavior of the nematode Caenorhabditis elegans, is regulated by IL2 neurons. Nat Neurosci. 13 nov 2011;15(1):107-12. doi:10.1038/nn.2975 PubMed PMID: 22081161.

48. Doroquez DB, Berciu C, Anderson JR, Sengupta P, Nicastro D. A high-resolution morphological and ultrastructural map of anterior sensory cilia and glia in Caenorhabditis elegans. Hobert O, éditeur. eLife. 25 mars 2014;3:e01948. doi:10.7554/eLife.01948

49. Cook SJ, Kalinski CA, Loer CM, Memar N, Majeed M, Stephen SR, et al. Comparative connectomics of two distantly related nematode species reveals patterns of nervous system evolution. Science. 31 juill 2025;389(6759):eadx2143. doi:10.1126/science.adx2143

50. Han Z, Lo WS, Lightfoot JW, Witte H, Sun S, Sommer RJ. Improving Transgenesis Efficiency and CRISPR-Associated Tools Through Codon Optimization and Native Intron Addition in Pristionchus Nematodes. Genetics. déc 2020;216(4):947-56. doi:10.1534/genetics.120.303785 PubMed PMID: 33060138; PubMed Central PMCID: PMC7768246.

51. Loer CM, Yim H, Geiger LT, Ramadan YH, Hampton MF, Bernal DV, et al. Identity and functions of monoaminergic neurons in the predatory nematode Pristionchus pacificus reveal nervous system conservation and divergence. bioRxiv. 16 oct 2025;2025.10.16.682888. doi:10.1101/2025.10.16.682888 PubMed PMID: 41279892; PubMed Central PMCID: PMC12632918.

52. Ramadan YH, Hobert O. Visualization of gene expression in Pristionchus pacificus with smFISH and in situ HCR. microPublication Biology. 6 juill 2024. doi:10.17912/micropub.biology.001274

53. Staum M, Abraham AC, Arbid R, Birari VS, Dominitz M, Rabinowitch I. Behavioral adjustment of C. elegans to mechanosensory loss requires intact mechanosensory neurons. PLOS Biology. juil 2024;22(7):e3002729. doi:10.1371/journal.pbio.3002729

54. Price MP, Thompson RJ, Eshcol JO, Wemmie JA, Benson CJ. Stomatin modulates gating of acid-sensing ion channels. J Biol Chem. 17 déc 2004;279(51):53886-91. doi:10.1074/jbc.M407708200 PubMed PMID: 15471860.

55. Liang X, Taylor M, Napier-Jameson R, Calovich-Benne C, Norris A. A Conserved Role for Stomatin Domain Genes in Olfactory Behavior. eNeuro. 21 mars 2023;10(3):ENEURO.0457-22.2023. doi:10.1523/ENEURO.0457-22.2023 PubMed PMID: 36858824; PubMed Central PMCID: PMC10035767.

56. Rödelsperger C, Meyer JM, Prabh N, Lanz C, Bemm F, Sommer RJ. Single-Molecule Sequencing Reveals the Chromosome-Scale Genomic Architecture of the Nematode Model Organism Pristionchus pacificus. Cell Rep. 17 oct 2017;21(3):834-44. doi:10.1016/j.celrep.2017.09.077 PubMed PMID: 29045848.

57. Sternberg PW, Van Auken K, Wang Q, Wright A, Yook K, Zarowiecki M, et al. WormBase 2024: status and transitioning to Alliance infrastructure. Genetics. 7 mai 2024;227(1):iyae050. doi:10.1093/genetics/iyae050 PubMed PMID: 38573366; PubMed Central PMCID: PMC11075546.

58. Madeira F, Madhusoodanan N, Lee J, Eusebi A, Niewielska A, Tivey ARN, et al. The EMBL-EBI Job Dispatcher sequence analysis tools framework in 2024. Nucleic Acids Res. 1 juill 2024;52(W1):W521-5. doi:10.1093/nar/gkae241 PubMed PMID: 38597606; PubMed Central PMCID: PMC11223882.

59. Nakayama K ichi, Ishita Y, Chihara T, Okumura M. Screening for CRISPR/Cas9-induced mutations using a co-injection marker in the nematode Pristionchus pacificus. Dev Genes Evol. 1 mai 2020;230(3):257-64. doi:10.1007/s00427-020-00651-y

60. Bonnard E, Liu J, Zjacic N, Alvarez L, Scholz M. Automatically tracking feeding behavior in populations of foraging C. elegans. Elife. 9 sept 2022;11:e77252. doi:10.7554/eLife.77252 PubMed PMID: 36083280; PubMed Central PMCID: PMC9462848.

61. R Core Team. R: A language and environment for statistical computing [Internet]. Vienna, Austria.: R Foundation for Statistical Computing; 2021. Disponible sur: https://www.R-project.org/.

