## Supplementary figures and images for "MEC-2/Stomatin is required for aversive behaviour but dispensable for prey detection in the predatory nematode *Pristionchus pacificus*"

### Supplemental Files

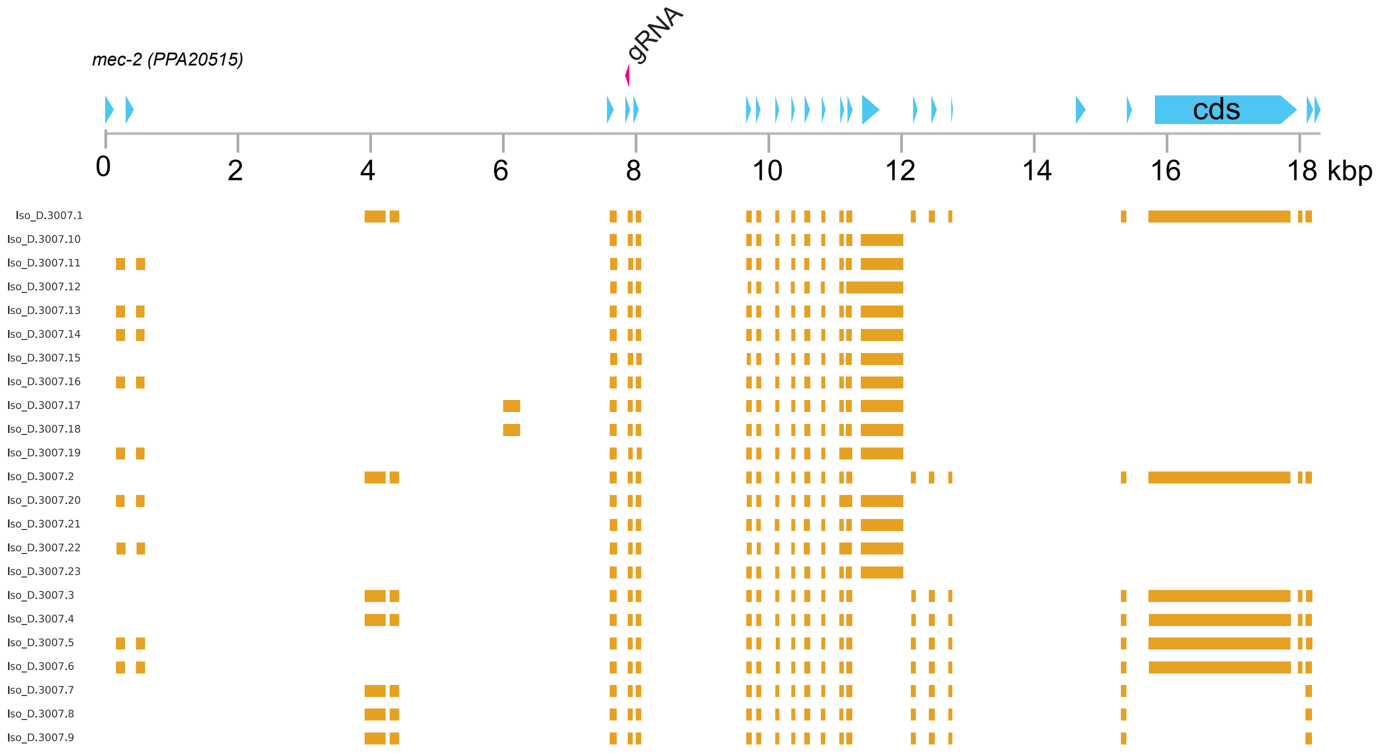

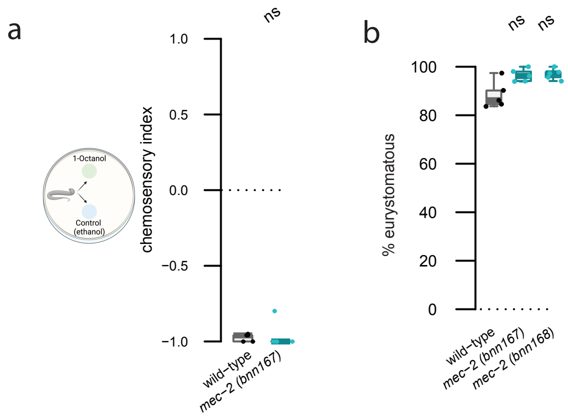
